# Rewinding the developmental tape shows how bears break a developmental rule

**DOI:** 10.1101/2024.02.29.582676

**Authors:** Otto E. Stenberg, Jacqueline E. Moustakas-Verho, Jukka Jernvall

## Abstract

Mammals have evolved a broad variety of dental morphologies. Nevertheless, the development of the mammalian dentition is considered highly conserved. Molar size proportions exemplify this as a system where small changes in shared developmental mechanisms yield a defined range of morphological outcomes. The Inhibitory Cascade (IC) model states that as molars develop in a sequence, the first developing anterior molars inhibit the development of subsequent posterior ones. The IC model thus predicts a trend of linear tooth size change along the molar row, as has been observed in a wide range of mammalian taxa with otherwise differing dental morphology. Perhaps the starkest exceptions to the IC rule are bears, in which the second molar is the largest and the third molar is disproportionally small. Here we sought to illuminate when and how during development the bear dentition falls of the IC prediction. We examined molar proportions in seven bear species. The results indicate that development of bear molars deviates from IC expectation already during patterning. Yet, during the earlier cap stage, size proportions of bear molars still seem to adhere to the IC model predictions. Overall, these analyses are suggestive that irrespective of the final outcome, the process of initial splitting of the molar-forming region into individual teeth is conserved and follows the IC rule.

## Introduction

Bear dentitions have received considerable research interest, undoubtedly due to factors such as bears being typically considered as apex predators. Different bear taxa also show remarkable diversity in their ecology making them highly suitable for comparative studies. Kurtén’s own work, starting on cave bear teeth, pioneered quantitative approaches that linked intra- and interspecies variation (Kurtén 1953). Some of his studies incorporated also perspectives relevant to developmental biology. These are evident already in Kurtén’s thesis work in which he examined the minimum size when teeth can develop, as also visualised integration of dentitions using correlation fields of tooth sizes (Kurtén 1953, see also Gomez-Robles and Polly 2012).

During the last 30 years, developmental biology studies have advanced to incorporate molecular evidence that has provided mechanistic insights into the processes regulating dental variation. One of these new insights is the interplay of signals regulating dental development along the anteroposterior sequence. This has been experimentally studied in molar teeth of the mouse, in which the presence of the first molar (m1) has been observed to inhibit the development of subsequent distal molars (Kavanagh et al. 2007). Cultivating mouse lower molars *ex vivo*, Kavanagh et al. (2007) microdissected explants, separating the developing m1 from the posterior tail that gives rise to the second (m2) and third (m3) molars. Comparing cut explants to the intact ones, Kavanagh et al. (2007) noted that the absence of the m1 significantly accelerated the initiation and growth of the m2 and the m3, elementally altering the morphology and size proportions of the resulting tooth row. This work implied that as molars are developing in succession, they are subject to cumulative effects of prior developmental events. In short, the state of the first tooth increasingly affects the development of subsequent ones. The authors introduced this proposed developmental ratchet as an inhibitory cascade (IC) model (Fig. 1a).

**Fig. 1.**
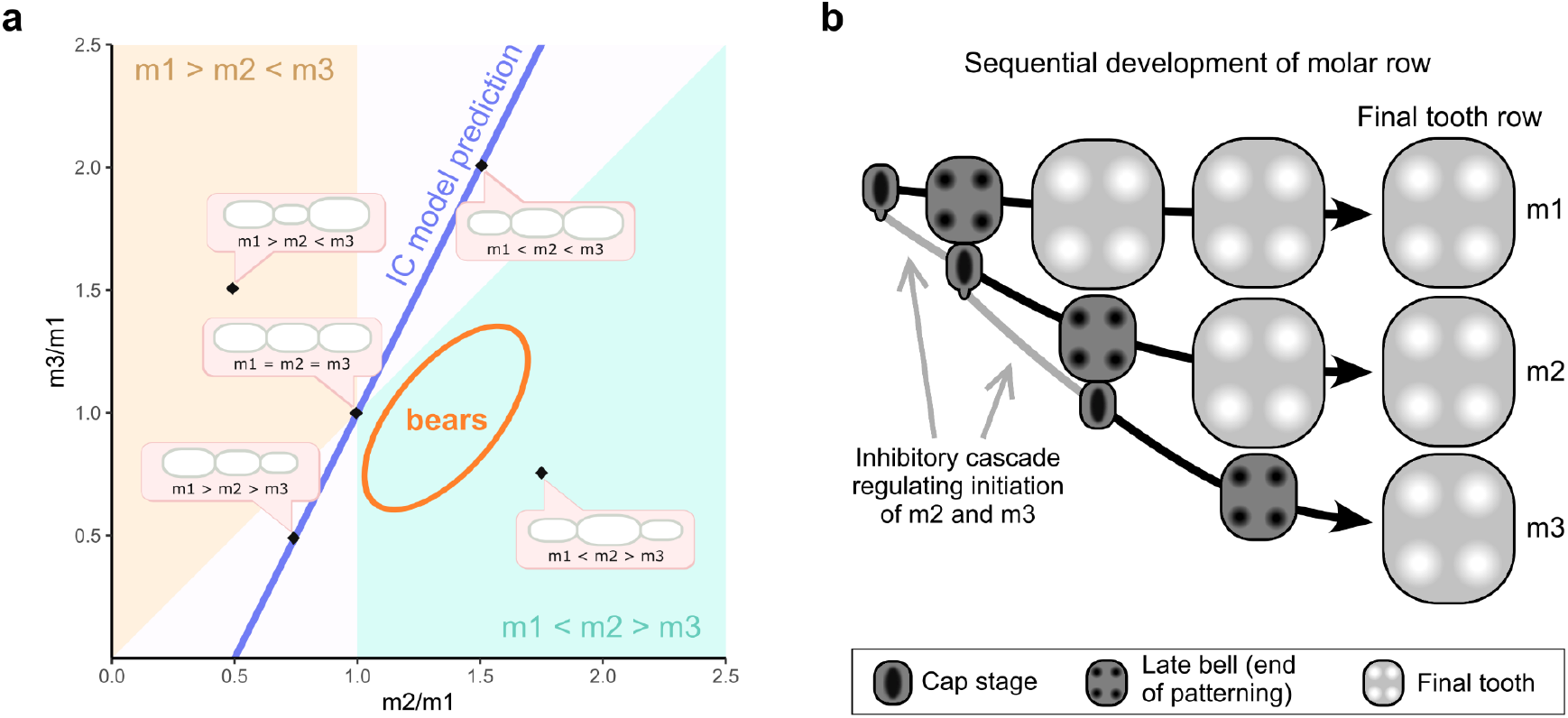
The IC model predicts linear tooth size change along the molar row. (**a**) Visualised in a morphospace of molar size ratios, the largest bear molar being the m2 is a departure from the expectations of the IC model. (**b**) Molars develop in a sequence, during which the previous tooth regulates initiation of the next one in an inhibitory cascade, producing a linear trend in tooth size along the molar row. In this study we analytically estimate the sizes of late bell and cap stage teeth.

Formally the IC model states that the proportions of molars are determined by the ratio mesenchymal activation and intermolar inhibition during development, with a balance between these signals yielding a row of molars equal in size (m1 = m2 = m3). Characterized as a cascade, the model by Kavanagh et al. (2007) predicts that any increase in inhibition has a cumulative effect on the size of distal molars, producing a pattern of decreasing size along tooth row (m1 > m2 > m3) that even extends to the complete loss of distal molars — as seen in felids. Accordingly, decreased inhibition should inversely affect development, increasing the size of distal molars (m1 < m2 < m3). These predictions by the IC model entail covariance between teeth; the known sizes of two molars can be used to predict the size and presence of the third one. Instead of calculating correlation matrices among fully formed teeth, here the starting point was logic derived from experimentation on developing mouse molars.

Because the experimentations on mouse teeth produced large, interspecies level changes in tooth proportions, Kavanagh et al. (2007) investigated molar sizes in murine rodents and reported molar size patterns that mostly adhered to the predictions of the IC model. Since then, a wide range of taxa has been used to evaluate the IC model on a macroevolutionary scale (Polly 2007; Asahara 2013; Bernal et al. 2013; Halliday & Goswami 2013; Schroer & Wood 2015; Evans et al. 2016; Carter & Worthington 2016, Selig et al. 2021). Overall, the IC model appears to explain much of the variation in several lineages, suggesting that evolutionary change in molar proportions has largely obeyed on shared developmental logic. This logic is not, however, the driving force of evolutionary change. Rather, it is ecology. Herbivory in mammals is linked to larger and more complex molars, and faunivory to smaller simpler molars (Evans et al. 2007; Selig et al. 2021). The inhibitory cascade can be hypothesized to have been the developmentally simplest solution to modifying tooth size along the jaw (Kavanagh et al. 2007). This mode of development intrinsically carries the side effect that the more distal, later developing molars show the largest changes.

Whereas the majority of observed mammalian taxa fit the predictions of the IC model to a high degree, a number of groups are known to deviate from the expectations of the IC model. One such group, already noted by Polly (2007) is bears. He observed that the dentitions of three bear species fell well outside the area of morphospace predicted by the IC model, exhibiting a pattern in which the largest molar is the second one (m1 < m2 > m3). This type of deviation has been found to be quite common (e.g. Bernal et al. 2013; Halliday and Goswami 2013), raising the question how these departures from the IC predictions occur during development.

To study the deviations from the IC predictions, first it is important to consider what is truly being measured to test the IC model. The predominant measures used are two-dimensional (2D) areas of the teeth, whether as simple length-width estimates of area or more accurate direct measurements of 2D area. Although three-dimensional measures of size can also be used, they provide comparable results to the 2D measures (Evans et al. 2016). What these different measures of size share is the fact that they are measuring the very end point of the developmental process generating each tooth (Fig. 1b). This includes the initiation, patterning, growth, matrix secretion, and mineralization of the dental hard tissues (Fig. 1b) — any of which may affect final tooth size. In addition to the interactions producing the inhibitory cascade dynamics, other genetic and hormonal factors affecting tooth development can influence the final outcome. Consequently, decomposing the inhibitory cascade from all the other factors regulating tooth size is undoubtedly more difficult at the lower taxonomic and population levels where the range of phenotypic variation is smaller (e.g., Roseman and Delezene 2019, Boughner et al. 2021, Bermúdez de Castro et al. 2021). Nevertheless, a recent analysis of a large dataset of primate dental variation suggests that IC aligns microevolution with macroevolution (Machado et al. 2023). Yet, clear exceptions to the IC such as the bear molars are of special interest as they can be used to examine when and how these deviations occur during development. To state this in more general terms, understanding exceptions to developmental models and rules should help us explain how these rules function in the first place.

Here we examine how the lower molars of different bear taxa obtain the m1 < m2 > m3 size pattern, and therefore how these dentitions fall below the line predicted by the IC model (Fig. 1a). Following the logic of sequential development of molars (Fig. 1b), we assume that most of the deviation is due to the last tooth of the cascade, the third molar, being too small, and focus on explaining deviation from the expected m1 < m2 < m3 pattern. We specifically ask when during development this deviation occurs. Fully formed bear molars are almost uniquely suitable for this particular question because their crowns are highly complex. Cusp features responsible for the surface complexity are principally formed during patterning stage of development, a process of which fully formed bear molars preserve a fine-grained proxy. Moreover, by analytically rewinding development, we estimate tooth sizes at the end of patterning stage and at the earlier cap stage when the patterning process is only just beginning (Fig. 1b). Taken together, by quantifying surface complexity and performing a developmental rewind, we can peer into earlier stages of development and ask whether the bear molars fall off the expected developmental trajectory before or after the patterning stage.

## Material and methods

### Sample

We sampled the lower molar dentition of seven bear species, encompassing all extant ursids but the spectacled bear (*Tremarctos ornatus*). Medians measurements were calculated for species with several sampled specimens (Table 1). The specimens originate from the collections of the Finnish Museum of Natural History (FMNH), Zoological Museum of the University of Copenhagen (ZMUC), and the Smithsonian National Museum of Natural History (USNM). Tooth rows were selected with the criteria of being complete and as unworn as available. The giant panda molar (*Ailuropoda melanoleuca*) row was a cast.

**Table 1.** The primary measurements from the sampled bear specimens. Medians are given on the first row of species with multiple specimens.

| Species, specimen | 2D area (mm<sup>2</sup>) |  |  | Complexity (OPCR) |  |  | Width (mm) |  |  |
| --- | --- | --- | --- | --- | --- | --- | --- | --- | --- |
|  | m1 | m2 | m3 | m1 | m2 | m3 | m1 | m2 | m3 |
| *Ailuropoda melanoleuca* |  |  |  |  |  |  |  |  |  |
| USNM 259400, right | 416.9 | 432.5 | 261.4 | 87.6 | 128.5 | 114.5 | 17.1 | 19.9 | 19.3 |
| *Helarctos malayanus* |  |  |  |  |  |  |  |  |  |
| FMNH 40_1976, left | 108.7 | 130.7 | 113.1 | 69.1 | 104.6 | 113.1 | 8.3 | 9.7 | 11.1 |
| *Melursus ursinus* |  |  |  |  |  |  |  |  |  |
| ZMUC 4290, left | 125.5 | 135.5 | 75.1 | 50.6 | 80.6 | 67.1 | 9.4 | 10.4 | 8.8 |
| *Ursus americanus*, median | 108.7 | 168.1 | 125.4 | 69.3 | 112.1 | 109.0 | 7.9 | 11.1 | 12.4 |
| FMNH 545 1960, left | 107.4 | 167.2 | 111.9 | 69.3 | 111.4 | 103.4 | 7.9 | 10.8 | 10.4 |
| FMNH 991, left | 108.7 | 168.1 | 125.4 | 71.6 | 115.9 | 143.1 | 7.9 | 11.1 | 12.4 |
| FMNH 996_1952, left | 136.1 | 206.3 | 173.0 | 60.4 | 112.1 | 109.0 | 8.7 | 11.9 | 13.1 |
| *Ursus arctos*, median | 173.9 | 248.2 | 202.1 | 78.9 | 124.9 | 111.4 | 10.0 | 12.9 | 13.6 |
| FMNH 1362, left | 187.9 | 251.0 | 203.5 | 80.0 | 128.5 | 94.5 | 10.4 | 12.9 | 14.6 |
| FMNH 1369, left | 171.5 | 245.3 | 192.3 | 71.0 | 125.8 | 111.8 | 9.9 | 12.9 | 13.0 |
| FMNH 22.679, left | 164.5 | 223.3 | 200.7 | 80.5 | 124.1 | 151.8 | 9.5 | 12.2 | 13.5 |
| FMNH 39733, left | 176.4 | 261.4 | 204.7 | 77.8 | 118.0 | 111.1 | 10.0 | 13.5 | 13.7 |
| *Ursus maritimus*, median | 157.8 | 194.6 | 160.9 | 63.6 | 93.0 | 112.6 | 8.7 | 11.2 | 12.4 |
| ZMUC 5365, left | 136.5 | 154.5 | 135.0 | 60.5 | 71.5 | 96.6 | 8.5 | 9.8 | 10.8 |
| ZMUC 5378, left | 163.5 | 209.4 | 160.9 | 63.6 | 93.0 | 112.6 | 9.1 | 11.2 | 12.9 |
| ZMUC 5620, left | 132.8 | 171.0 | 93.4 | 60.4 | 99.3 | 86.6 | 8.2 | 10.8 | 9.8 |
| FMNH 2342_201, left | 168.5 | 222.3 | 161.1 | 65.0 | 106.9 | 131.6 | 8.9 | 11.8 | 12.5 |
| ZMUC 5472, right | 157.8 | 194.6 | 163.2 | 67.3 | 85.3 | 114.4 | 8.7 | 11.4 | 12.4 |
| *Ursus thibetanus*, median | 151.7 | 191.2 | 151.9 | 70.3 | 98.8 | 83.2 | 8.4 | 11.4 | 11.8 |
| FMNH 1.962, left | 152.9 | 210.5 | 190.3 | 66.6 | 99.3 | 100.0 | 8.5 | 11.7 | 13.0 |
| ZMUC 603, right | 150.5 | 171.8 | 113.5 | 73.9 | 98.4 | 66.4 | 8.3 | 11.1 | 10.6 |

### 3D scanning and mesh preparation

Lower molar rows were as surface scans using a Planmeca PlanScan intraoral dentistry scanner (PlanMeca, Helsinki, Finland). The surfaces were prepared for analysis in MeshLab (v. 2020.07, Cignoni et al. 2008) as described in Christensen et al. (2023) with the following modifications: no smoothing was deemed necessary for surface scans and the quadric edge collapse decimation target face count was set to 20,000. After this, molars were separated. As the bear m3 is generally tilted more lingually than the other molars, each m3 was manually realigned to have the occlusal surface aligned horizontally.

### Measurements and estimation of tooth size during development

OPCR was measured and OPC maps produced following Christensen et al. (2023). In addition, the two-dimensional projection area of each tooth was measured in mm^2^ utilizing Morphotester’s (v. 1.1.2, Winchester 2016) RFI functionality. Maximum width was measured for each molar in mm. A feature density was obtained as a ratio of complexity to area (OPCR/mm^2^) for each tooth. Area, complexity, and feature density proportions of all molar tooth rows were calculated (m2/m1, m3/m1). For species with several sampled specimens, medians of these ratios were calculated and used in the figures.

For each species, we estimated tooth area at two points of development. Utilizing equations from the work of Christensen et al. (2023), we used final tooth width and area to estimate developing molar size (mm^2^) at the end of patterning (*A*_pat_, Eq. 1) and at the onset of patterning (*A*_cap_, Eq. 2). These stages correspond to the late bell and cap stages of crown formation, respectively (Fig. 1b). For *A*_pat_ and *A*_cap_ we write

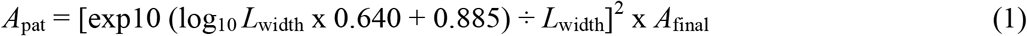

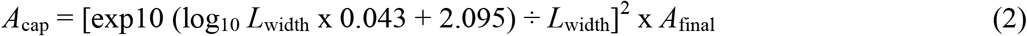

where *L*_width_ denote to the maximum width and *A*_final_ to the area of the fully formed tooth, respectively. Units are in micrometres. Note that the original equations in Christensen et al. (2023) are for the widths of the developing teeth at the cap and late bell stage. Here we assume that the width-length proportions of the crowns are roughly constant from the cap stage onwards. This is supported by the observation that length differences between developing teeth appear first (Christensen et al. 2023). Thus, the cap stage when the forming grown base is established by lateral expansion of the cervical loops, is the earliest stage when the crown proportions can be observed. In addition, although the original equations were derived from data on individual teeth from different species, here we apply the equations also to the teeth in the same tooth row.

## Results

### Both proportions in molar size and complexity show bears to be outliers of the IC rule

A simple plot of molar sizes along the tooth row makes it immediately apparent that the second molar is largest in size while the third molar is smaller, (m1 < m2 > m3) in all sampled bear dentitions (Table 1, Fig. 3a). This pattern is inconsistent with the IC model of linear change from tooth to tooth as the m3 is much smaller than would be expected from the m2/m1 ratio (Fig. 3b). In the case of the *Ailuropoda* and *Melursu*s, the m3 is by far the smallest molar in the tooth row (Fig. 2, Fig. 3b), making these species fall further down in the IC morphospace (Fig. 3b, the mean distance from the IC line is 0.257 and the range is 0.173 to 0.433).

**Fig. 2.**
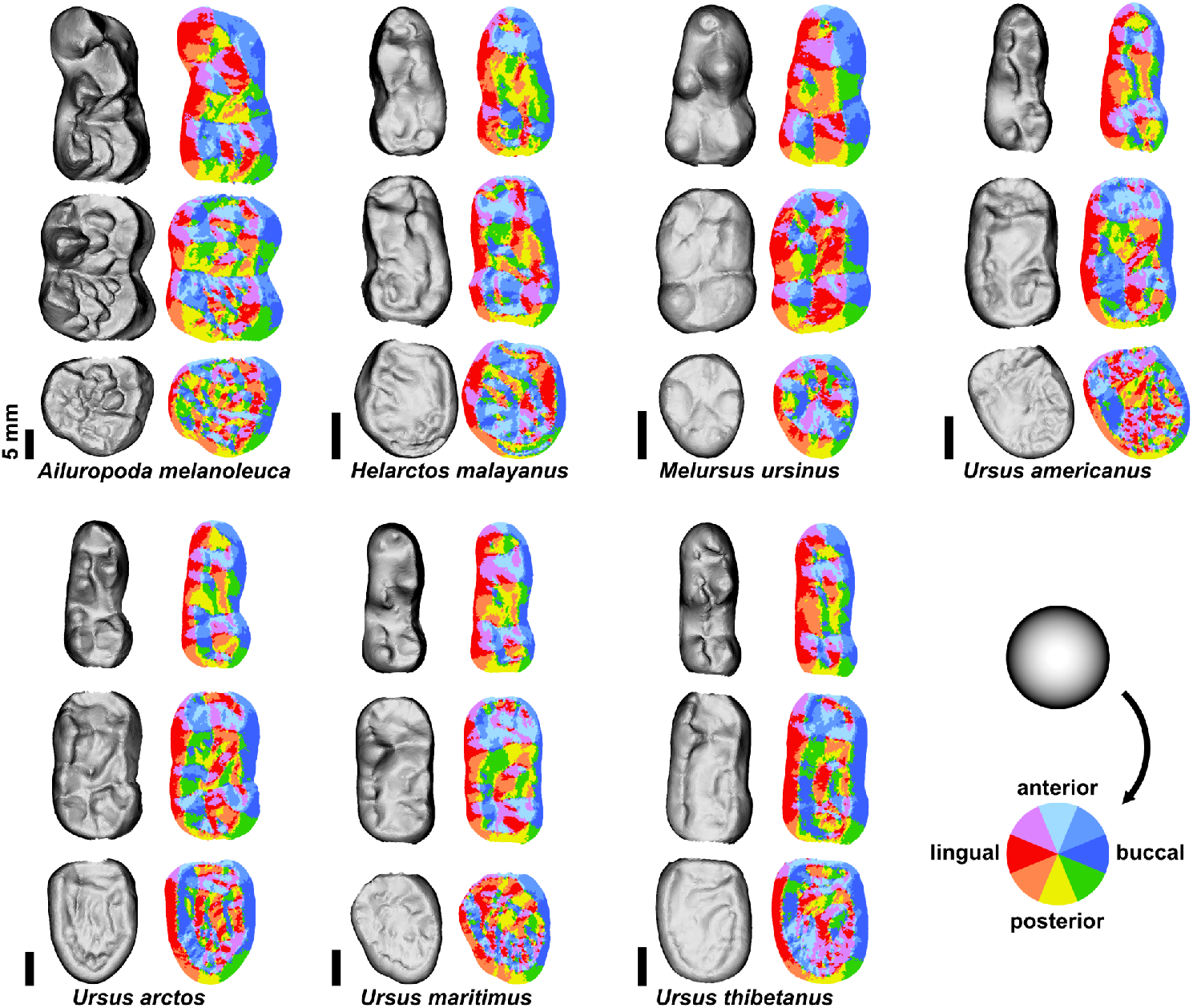
Three-dimensional surfaces and OPC maps illustrate differences in dental complexity across the seven bear species sampled. For species with several specimens, a representative tooth row is shown. Left side molar rows were mirrored for consistency. Scale bars, 5 mm.

**Fig. 3.**
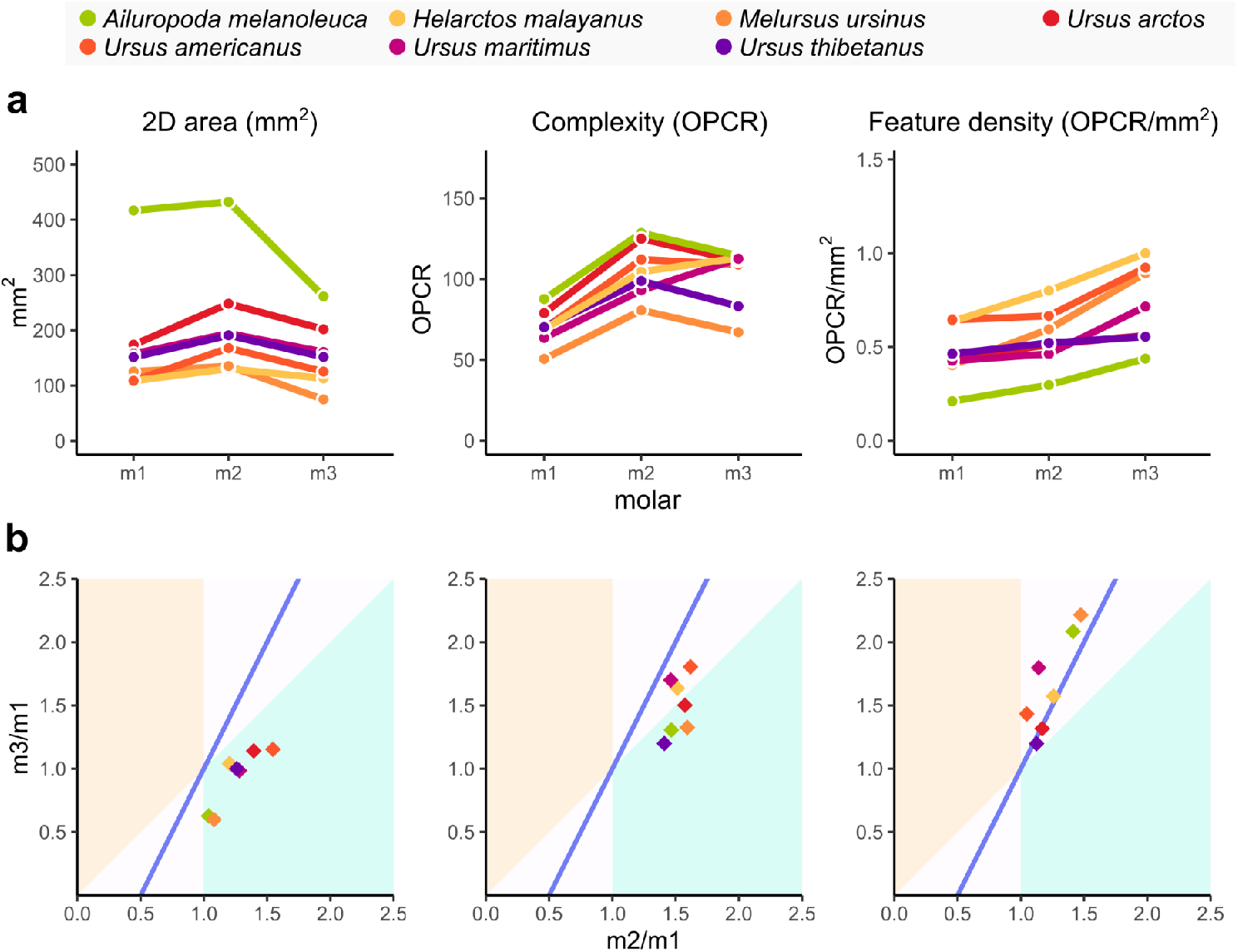
Molar area, complexity, and feature density along the molar row. Molar area and complexity do not change linearly along the molar row, but feature density is highest in the m3 as can be observed from (**a**) tooth sizes, and (**b**) tooth proportions plotted in the IC morphospace. In (**b**), the blue line denotes the IC prediction and coloured regions either m1 > m2 < m3 above, and m1 < m2 > m3 below the IC line, respectively.

Next, we examined the complexity of the same molars using the orientation patch count rotated (OPCR) measure. The pattern of m2 being the largest tooth holds for the OPCR values in most of the species (Fig. 3a). *U. maritimus* and *Helarctos* are exception in that their m3s have the largest OPCR values, and they are also closest to the IC line (Fig. 3a,b). Overall, apart from *U. maritimus*, the species appear to still fall below IC line (the mean distance from the IC line is 0.260 and the range is 0.068 to 0.384). Because the offset from the IC line remains roughly similar between the size and complexity, we also illustrated the feature density, calculated as the ratio of complexity to area. The density of features increased along the tooth row (m1 < m2 < m3, Fig. 3a), and the sample straddled along or above the IC line (Fig. 3b, the mean distance from the IC line is 0.081 and the range is 0.002 to 0.163). This result is indicative of both the patterning and final growth stages being roughly equally affected, pointing towards an early developmental divergence from the IC.

### Rewinding development reveals the beginning of the deviation from the IC

Using two empirically derived equations (Christensen et al. 2023) we estimated the sizes of developing teeth at the late bell stage, which corresponds to the end of patterning stage, and at the earlier cap stage. Crown morphogenesis starts at the cap stage when the cervical loops begin to grow laterally to the primary enamel knot, and thus these two estimates, together with the final tooth size, bracket the whole crown morphogenesis (Fig. 1b). The resulting estimates show that late bell stage molars are closer to the IC line, but still fall below it (Fig. 4a,b, the mean distance from the IC line is 0.163 and the range is 0.108 to 0.275). Compared to the fully formed teeth, the relative size of the m2 is smaller and more equal in size to m1, or even smaller as in *Ailuropoda*. (Fig. 4a). Calculating the cap stage size estimates reveals a trend of relatively linear reduction in sizes along the tooth row (m1 > m2 > m3, Fig. 4a). This in turn means that the cap stage sizes appear to agree with the IC prediction (Fig. 4b, the mean distance from the IC line is 0.046 and the range is 0.002 to 0.095). Analytically, the change in tooth proportions in the developmental rewind is explained by differences in tooth lengths and widths. The size proportion estimates for the three cap stage molars converge around 1.0, 0.75, and 0.5 for the m1, m2, and m3, respectively. To the extent these estimations reflect the actual developmental process, this point on the IC line equals activator/inhibitor ratio of 0.75 in the original inhibitory cascade formula (Kavanagh et al. 2007).

**Fig. 4.**
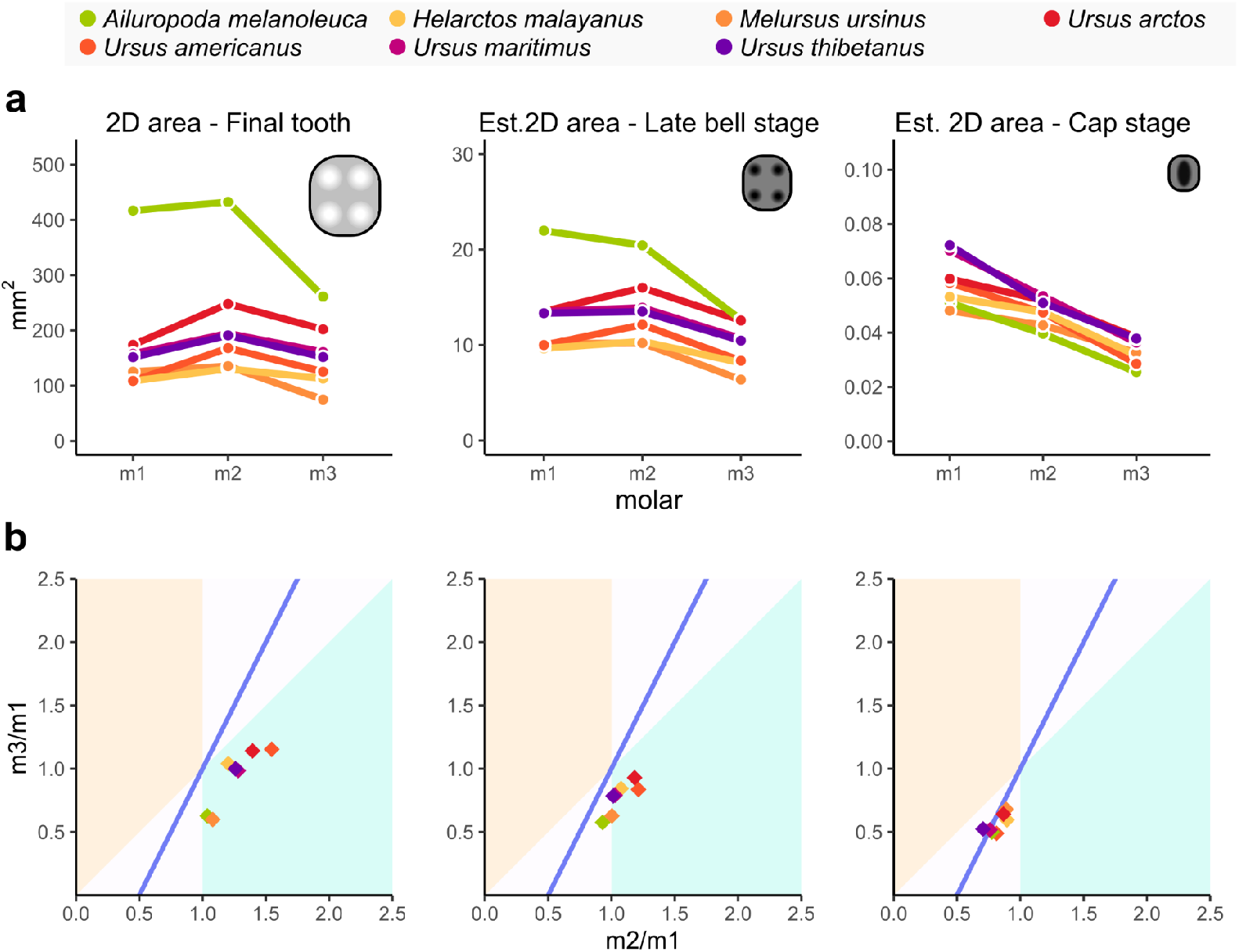
Tooth sizes (**a**) and their position in IC morphospace (**b**) at three points of development. Estimates of size at the late bell and the cap stage are based on an analytical rewind of the development (Fig. 1b). Whereas the late bell stage estimates remain below the IC line, the cap stage estimates appear to follow the IC prediction of linear change is size between teeth. In (**b**), the blue line denotes the IC prediction and coloured regions either m1 > m2 < m3 above, and m1 < m2 > m3 below the IC line, respectively.

## Discussion

The opportunity to study mammalian tooth development in action is limited to a handful of species. A common feature of these species that can be investigated experimentally is their small size. Despite its highly derived dentition, the mouse has remained the species that has contributed most to our understanding of tooth development. In this context bears make poor model organisms. But from a standpoint of evolutionary diversity, bears provide prime examples to test on why generalizations made from mouse tooth development sometimes fall short.

The dental inhibitory cascade is a proposed, seemingly plesiomorphic system of development, which describes how a balance of inhibition and activation drives the sequential development of molars, resulting in linear patterns of tooth size along the molar row (Kavanagh et al. 2007). Though most of Mammalia largely fit these predictions of molar proportions, bears are a noted exception to this. Moreover, the mode of departure from the IC prediction (m1 < m2 > m3) is commonly observed in other mammalian groups such as primates, and also at lower taxonomic levels (Bernal et al. 2013; Halliday and Goswami 2013). Explanations on how bears escape the IC may thus apply more broadly.

Here we analytically rewound tooth development in bears by integrating recent advances in developmental theory with computational methods for quantifying tooth shape. Tooth shape is chiefly established by two partly concurrent developmental processes: patterning and growth (Fig. 1b). Measuring crown complexity along the molar row in the context of the IC model provided a proxy for the patterning process preceding the final growth in size. Selig et al. (2021) showed that in primates and related groups the complexity of tooth crowns generally increases from m1 to m3, and shows better fit to the IC than the corresponding molar sizes. The size pattern was more alike the one in bears. The more intricate bear dentitions paint a somewhat different picture: similar to the molar size ratios, the proportions of complexity in bears remain below the IC prediction (Fig. 3). This may imply that whereas in primates the development follows the IC prediction through the patterning, the bears are divergent already before patterning. This is supported when shape and size are considered together. A ratio of complexity to area — summarizing how fine-grained the morphological detail of surface is — shows a closer match with the IC line (Fig. 3). Though the m2 displayed higher feature density than the m1, the m3 far surpassed both in this aspect. A way to interpret this result is that both patterning and the growth after patterning are affected equally because the number of cusps and crown features in general are ultimately limited by the size of the growing tooth.

Our second approach to rewind tooth development took advantage of the empirical discovery that the size when the patterning is completed, and the size when the patterning begins, show strict scaling relationships with the final tooth size (Christensen et al. 2023). The mechanistic explanation for these relationships is the integration of patterning and growth processes by insulin-like growth factor (IGF) pathway. Whereas in larger teeth the patterning scales so that it happens at a larger size, the initiation of patterning is relatively size invariant (Christensen et al. 2023). We found that the estimated late bell size, when the patterning is completed, is closer to the IC line, but still below it (Fig. 4). This fits the result on the complexity and suggests that the relatively small size of bear m3, for example, is not a result of a developmental arrest late during development. Rather, the departure from the IC appears to involve most parts of the crown formation process.

The discovery that the estimated cap stage teeth are on the IC line is intriguing as it indicates that the initial splitting of the tooth-forming region follows the IC rule (Fig. 4). The cap stage is also closest in time to the actual inhibitory cascade tested experimentally by Kavanagh et al. (2007). That is, the microdissection separating the developing m1 from the posterior tail was done when the m1 was at the cap stage. While our results on bear teeth should be considered preliminary, it is perhaps a plausible hypothesis that the IC mode of tooth formation is the conserved, or plesiomorphic (Halliday & Goswami 2013) feature of all mammals. And whereas bears are one group that has diverged greatly from the IC, at the onset of the crown formation they still follow the rule. These kinds of analyses may also help to classify and uncover different mechanisms that produce the divergent patterns. One important aspect related to the cascading mode of development is that the late bell and cap stage molars are not at the same developmental stage at the same time (Fig. 1b). This means that the plots placing the developing teeth into the IC morphospace manifest the process of development, not a static state of morphology (Fig. 4). In this context it is also useful to keep in mind that molars in a fully formed tooth row, while typically analysed as a static representation of anatomy, actually reach their final sizes at different points in time (Fig. 1b).

In conclusion, analyses of morphology that utilize theories derived from developmental processes can be used to both test these theories, and to provide new insight into the kinds of developmental changes producing phenotypic diversity. In the case of mammalian teeth and the IC, it will be interesting to decipher the number of ways in which this rule can be broken — both from the standpoints of evolution and development.

## Acknowledgments

We thank the members of Jernvall lab for comments and discussions on this work and J. Granroth (Finnish Museum of Natural History, Helsinki) and E. Lorenzen, (Zoological Museum of the University of Copenhagen) for access and help with museum collections. This study was supported by the Academy of Finland (J.J.), SYNTHESYS Grant DK-TAF-7064 (J.E.M.-V.), and Finnish Doctoral Programme in Oral Sciences (O.E.S.).

